# Physiology and structure of pathogenic *Escherichia coli* pOmpT reveal two substrate-binding sites

**DOI:** 10.1101/2022.06.17.496525

**Authors:** Juanhua Liu, Tingting Ran, Luyao Jiang, Qingqing Gao, Changchao Huan, Hang Wang, Weiwu Wang, Song Gao, Xiufan Liu

**Author notes:** Contributed equally to this article (Juanhua Liu, Tingting Ran, Luyao Jiang).

## Abstract

The pathogenicity of bacteria can be achieved by cleaving antimicrobial peptides (AMPs) through the outer membrane protease family (Omptins) to evade or resist the host’s innate immune response. The OmpT-like proteins in *Escherichia coli* are members of Omptins, which have highly conserved proteolytic activity. Here, sequence alignment and physiological studies have determined that pOmpT is a virulence factor with atypical proteolytic activity. Comparing pOmpT with cOmpT in terms of structure, proteolytic activity and target substrates, it is found that Asp^267^ and Ser^276^ of cOmpT are substrate-binding sites and help its catalytic center to cleave substrates (protamine,synthetic peptide or RNase 7). However, special structural features and the nature of residues Ser^267^ and Thr^276^ of pOmpT caused the inability of cleavage protamine, but allows it to specifically cleave the human AMP RNase 7. It is suggested that two sites are related with the substrate specificity. In short, we found that pOmpT presents a new structural basis for the specific recognition of substrates, and providing new clues for the development of antimicrobial drugs.

## Introduction

*Escherichia coli* (*E. coli*) is a common commensal bacterium in the gastrointestinal tract of mammals and birds. However, it is also a multifunctional pathogenic bacterium associated with a variety of intestinal and extraintestinal infections, causing a variety of life-threatening diseases in humans and animals. At present, on account of the complex diversity of virulence factors of pathogenic *E. coli*, bacterial antibiotic resistance and ineffective vaccine control pose a potential threat to the healthy and sustainable development of the poultry industry and public health. Therefore, there is an urgent need to seek more effective prevention and treatment methods.

Antimicrobial peptides (AMPs) is one of the many initial risks faced by bacterial pathogens infecting the host, and is an important part of the innate immune system (Samantha & Le, 2012). These small (20-50 amino acids), cationic and amphiphilic peptides are mainly released from the host’s epithelial cells and neutrophils (Zasloff & Michael, 2002; Hancock & Sahl, 2006; Gallo & Hooper, 2012). They can bind to anionic bacterial membranes and form pores, and ultimately bacteria are killed due to cell lysis (Piers & Hancock, 1994; Zhang *et al*, 2001; Brogden, 2005). In addition to bactericidal activity, AMPs can recruit host’s immune cells to the site of infection to induce a broad range of immunomodulatory functions to control bacterial infection (Hilchie *et al*, 2013).

Despite the importance of AMPs in host response, bacteria have evolved several mechanisms to resist the actions of AMPs, including the use of LPS modifications, efflux pumps, capsules, and proteases (Samantha & Le, 2012). As OM proteases with proteolytic activity, omptins have attracted extensive attention. Omptins are a unique family of outer membrane proteases commonly found in a variety of Gram-negative bacteria in Enterobacteriaceae. They affect bacterial virulence by processing or degrading a variety of host and bacterial proteins (Grodberg & Dunn, 1988; Sodeinde *et al*, 1992; Kukkonen & Korhonen, 2004; Lathem *et al*, 2007; Haiko *et al*, 2009; Le Sage *et al*, 2009; Franco *et al*, 2011; Thomassin *et al*, 2012; Brannon *et al*, 2013; Korhonen *et al*, 2013; Caulfield *et al*, 2014). At present, the reported Omptins mainly include OmpT, OmpP and ArlC of *E. coli*, Pla of *Yersinia pestis*, PgtE of *Salmonella enteritidis*, IcsP of *Shigella flexneri* and CroP of *Citrobacter rodentium*, and they share high amino acid sequence identity (45-80%) with highly conserved active sites (Grodberg & Dunn, 1988; Grodberg & Dunn, 1989; Sodeinde & Goguen, 1989; Vandeputte-Rutten *et al*, 2001; Le Sage *et al*, 2009; Egile *et al*, 2010; Desloges *et al*, 2019). OmpT, encoded by the nucleoid of *E. coli* and firstly structured, is a 37 kDa outer membrane protein with a hollow β barrel structure and conserved active sites facing the extracellular environment (Kramer *et al*, 2001; Vandeputte-Rutten *et al*, 2001). Previous studies on omptin inhibition reported that Zn^2+^, Cu^2+^, and benzamidine are able to inhibit OmpT activity (Sugimura & Higashi, 1988; Sugimura & Nishihara, 1988; Yam *et al*, 2001), and the serine protease inhibitors aprotinin and ulinastatin interfere with the activity of OmpT (Gill *et al*, 2000; Hui *et al*, 2010). In addition, OmpT’s highly conserved structural features and the residues’s nature of the active site determine the specificity of cleavage substrates and can preferentially cleave substrates at dibasic motifs (Dekker *et al*, 2001; McCarter *et al*, 2004; Varadarajan *et al*, 2005). Studies have reported that OmpT in urinary pathogenic *Escherichia coli* (UPEC) can help bacteria survive in the host by cleaving the antimicrobial peptides protamine P1 and cathelicidin LL-37 secreted by human urethral epithelial cells (Stathopoulos, 1998; Stumpe *et al*, 1998; Brannon *et al*, 2013; Xiao *et al*, 2015). Meanwhile, LL-37 is cleaved and inactivated to different extents by OmpT present in enterohemorrhagic *E. coli* (EHEC) and enteropathogenic *E. coli* (EPEC) (Thomassin *et al*, 2012; Thomassin *et al*, 2012). The plasmid encoding OmpT-like protease OmpP found in urinary tract infections (UTIs) isolates has been shown to cleave the AMP protamine and ArlC is associated with AMP resistance (such as human AMP RNase 7), thereby helping bacteria survive (Hwang *et al*, 2007; Mcphee *et al*, 2014; Desloges *et al*, 2019).

Omptins are potential targets for antimicrobial and vaccine development as a key virulence factor that plays a core role in the host-pathogen interface, so it has research potential. In fact, for most *E. coli*, OmpT is only encoded by the *ompT* gene on the nucleoid. Previous researches mainly focused on the proteolytic activity and the pathogenicity of OmpT located on the nucleoid. We found that avian pathogenic *E. coli* (APEC) E058 strain, one of the extraintestinal pathogenic *E. coli*, has both OmpT (cOmpT) encoded by gene (c*ompT*) on the nucleoid and OmpT (pOmpT) encoded by *ColV* plasmid gene (p*ompT*), and both are involved in the pathogenicity of APEC on the host (Chen, 2016; Liu *et al*, 2019). Although cOmpT is extensively studied, the research on pOmpT is relatively lacking. Here, we aim to explore the structural characteristics of pOmpT and the special role it plays in physiology compared with cOmpT. The in-depth study on proteases like OmpT is conducive to design new enzyme-resistant antimicrobial peptides, and also provides a possibility for the exploration of new anti-infection approaches and the screening of new drug targets.

## Results

### pOmpT is an outer membrane protein like cOmpT

Sequence alignment showed that the primary structure of APEC E058 pOmpT is 76.45% identical to cOmpT of UPEC and E058 (Fig 1C), and pOmpT of E058 shares 226/297 amino acid sequence identity with cOmpT, in contrast, pOmpT of E058 shares 297/297 with the ArlC encoded by plasmids in UPEC, suggesting that these plasmids encoded OmpTs may have a same ancestor. RT-PCR result showed that both c*ompT* and p*ompT* genes can be transcribed normally and were 954 bp in the APEC E058 strain (Fig 1A). Although the pOmpT shares less than 80% sequence identity with cOmpT, the key residues of the OmpT protease between E058 pOmpT and cOmpT are exactly the same, including the reported catalytic sites and the active sites. However, there are two residues in the LPS-binding sites of E058 strain pOmpT different from cOmpT strains (Fig 1C). The sequence alignment suggests that E058 cOmpT is a typical outer membrane proteases (Grodberg & Dunn, 1988). It is unclear whether pOmpT is same as cOmpT and located in outer membrane, albeit it shares high sequence identity with cOmpTs and also transcripted in E058. To check the location of pOmpT, the outer membrane proteins of wild-type strain E058 and c*ompT*/p*ompT* single/double-gene deletion mutants were extracted, and detected by western blotting using anti-OmpT monoclonal antibody. The results showed that a single band can be detected in the E058 c*ompT*/p*ompT* single-gene deletion mutants, and the size of the bands is different in both strains suggesting that two different proteins expressed in these two deletion mutatnts (Fig 1B). Meanwhile, two bands in the wild-type strain E058 can be detected, and the position of the bands is consistent with that observed in the E058 c*ompT*/p*ompT* single-gene deletion mutants, respectively (Fig 1B). No bands were detected in the c*ompT*/p*ompT* double-gene deletion mutants (Fig 1B). Above results indicated that both pOmpT and cOmpT are expressed on the outer membrane of E058.

**Figure 1.**
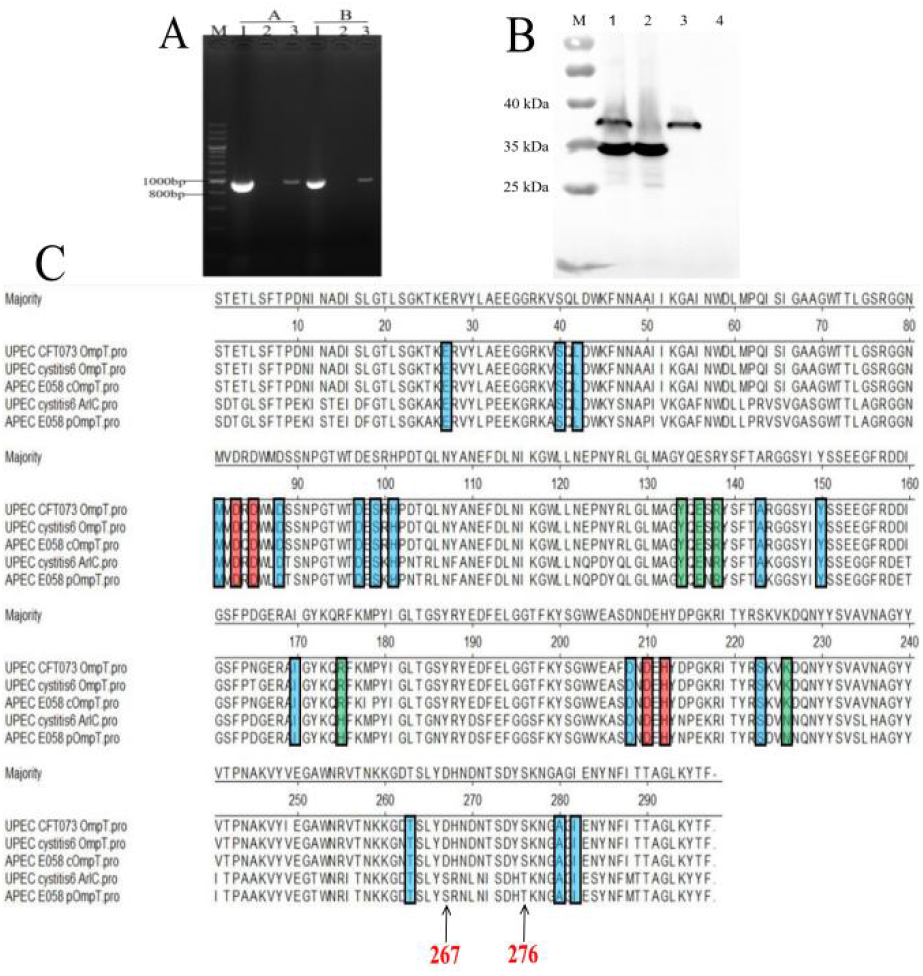
pOmpT is an outer membrane protein. A RT-PCR analysis of c*ompT* and p*ompT* genes in APEC E058 strain. Lane A: c*ompT* gene; Lane B: p*ompT* gene; Lane 1: genomic DNA from E058; Lane 2: total RNA from E058; Lane 3: cDNA derived from the total RNA of E058. 200 bp DNA marker (TaKaRa) was used as the molecular size standard (lane M). B The extracted total outer membrane proteins were also analyzed with anti-OmpT antibodies. Lane 1: Total outer membrane protein extracted from strain E058; Lane 2: Total outer membrane protein extracted from strain E058Δc*ompT*; Lane 3: Total outer membrane protein extracted from strain E058Δp*ompT*; Lane 4: Total outer membrane protein extracted from strain E058Δc*ompT*Δp*ompT*; PageRuler™ prestained protein ladder (Thermo Fisher scientific, USA) was used as the molecular size standard (lane M). C Alignment of the amino acid sequences between OmpT-like protein from different strains. The sequence of the nucleoid-encoded OmpT (E058 cOmpT) and the plasmid-encoded OmpT (E058 pOmpT) in the APEC E058 strain was determined by ourselves; The amino acid sequence of OmpT in UPEC CFT073 is from GenBank No. CP051263.1; The amino acid sequence of OmpT in UPEC isolate cystitis 6 comes from GenBank No. CP041302.1; The amino acid sequence of ArlC in UPEC isolate cystitis 6 is from GenBank No. CP041301.1. In the figure, the residues marked in the red box are the catalytic residues of OmpT outer membrane protease, the residues marked in the blue box are the enzyme active sites of OmpT and the green box marks the amino acid as the binding site of OmpT and LPS.

### pOmpT cannot resist protamine

Protamine cleavage is one of the important roles for cOmpT for host resistance (Stathopoulos, 1998; Stumpe *et al*, 1998; Brannon *et al*, 2013; Xiao *et al*, 2015). In order to understand the role of pOmpT in resisting the host, the growth kinetics were carried out for wild-type strain E058, c*ompT*/p*ompT* single/double-gene deletion strains and its complementation strains incubated with protamine. The results showed that compared with the wild-type strain, there was no significant difference in the growth of the strains E058Δp*ompT* and ReE058Δc*ompT-*cc*ompT* when cOmpT was present in the strains (*P*>0.05), while its growth ability was significantly reduced when only pOmpT was present in the strains E058Δc*ompT* and ReE058Δc*ompT-*pp*ompT*) (*P*<0.01) (**Fig 2A**). The same phenomenon has been verified in the recombinant expression bacteria of c*ompT*/p*ompT* genes (**Fig 2B**), which is a cue that cOmpT can resist protamine, but pOmpT cannot.

**Figure 2.**
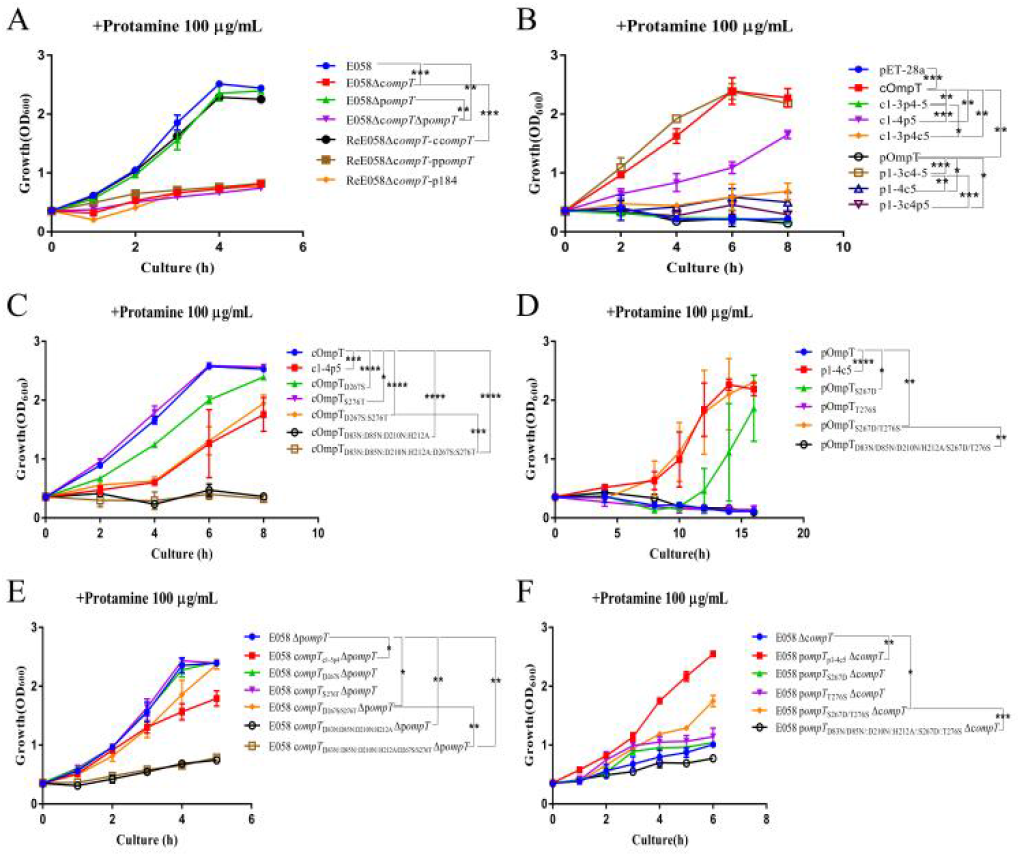
The kinetics of bacterial growth under protamine treatment. A Growth profiles of APEC wild-type strain E058, c*ompT*/p*ompT* single/double-gene deletion strains and its complementation strains; B Growth profiles of recombinant expression bacteria expressing cOmpT, pOmpT and the chimeric protein with the interchanged different loops between cOmpT and pOmpT; C Growth profiles of recombinant expression bacteria expressing site-directed mutagenesis of cOmpT; D Growth profiles of recombinant expression bacteria expressing site-directed mutagenesis of pOmpT; E Growth profiles of p*ompT* gene deletion strains expressing site-directed mutagenesis of cOmpT; F Growth profiles of c*ompT* gene deletion strains expressing site-directed mutagenesis of pOmpT. Data information: lines (A-F) represent mean values with standard error of the mean (n=5 cultures for each genotype). Statistical significance was determinated using the two-way ANOVA. Differences with p-values < 0.05 were considered as statistically significant. *: *p* < 0.05; **: *p* < 0.01; ***: *p* < 0.001; ****: *p* < 0.0001.

### Crystal structure of the pOmpT

pOmpT used for crystallization contains a mutation K217G to abolish autoproteolytic activity. pOmpT was crystallized in the three different space groups P3221 with unit cell dimensions a = b = 93.765 and c =150.344 Å, and P3121 with unit cell dimensions a = b =92.375 and c =74.434 Å and the best crystals were diffracted to resolutions of 2.95 Å (Fig 3A and B). The structure was solved by molecular replacement using cOmpT structure as template (Protein Data Bank with accession No.1I78). The asymmetric unit contains two monomers and one monomer, respectively (Fig 3A and B). Since the structures of two different space groups are almost identical with rmsd of 0.54-0.58 Å, we used the best resolution structure to present here. pOmpT forms a typical OmpT fold, a vase-shaped antiparallel β-barrel. The overall structure of pOmpT consists of all beta strands, no helices in the structure. The longest dimension of the barrel is around 70 Å. Among these strands, only the first strand is intact, all other beta strands are discontinues and formed by two beta strands (Fig 3A). Six of them (2, 3, 7, 8, 9 and 10) are broken into two strands at the surface region of the out membrane. Interestingly, a relative long region of the protein from residues 259 to 281 is not resolved (no electron density was observed for this region), the corresponding region in cOmpT contains two short beta strands (Fig 3D), the missing of this region suggests that the conformation of this region is flexible in pOmpT. In our structure, due to the crystal packing, the position of the missing region is occupied by two strands (the outside portion of seventh and eighth strands) of the symmetric molecule (Fig 3C and E). Protamine can completely inhibit the growth of *E. coli* BL21(DE3) strain, enzyme cleavage of protamine would inactivate its anti-bacteria function. Pretreatment of protamine with refolded cOmpT abolished the growth inhibition of *E. coli* BL21(DE3) strain by protamine, indicating that protamine was destroyed by cOmpT. Whereas pretreatment of protamine with pOmpT did not abolish the growth inhibition by protamine indicating that protamine was not inactivated by pOmpT. These results are consistent with the in vivo expression results, and also indicating that the refolded OmpTs are similar to the in vivo expressed protein and have similar function (Table 1). It is a cue that conformational flexibility of the missing of the region in pOmpT and makes the upper rim of the vase structure incomplete causing the substrate binding pocket dramatically different from cOmpT. The difference in substrate binding pocket may contribute to the activity loss to protamine of pOmpT comparison to cOmpT (Fig 2A and B; Table 1).

**Table 1.**
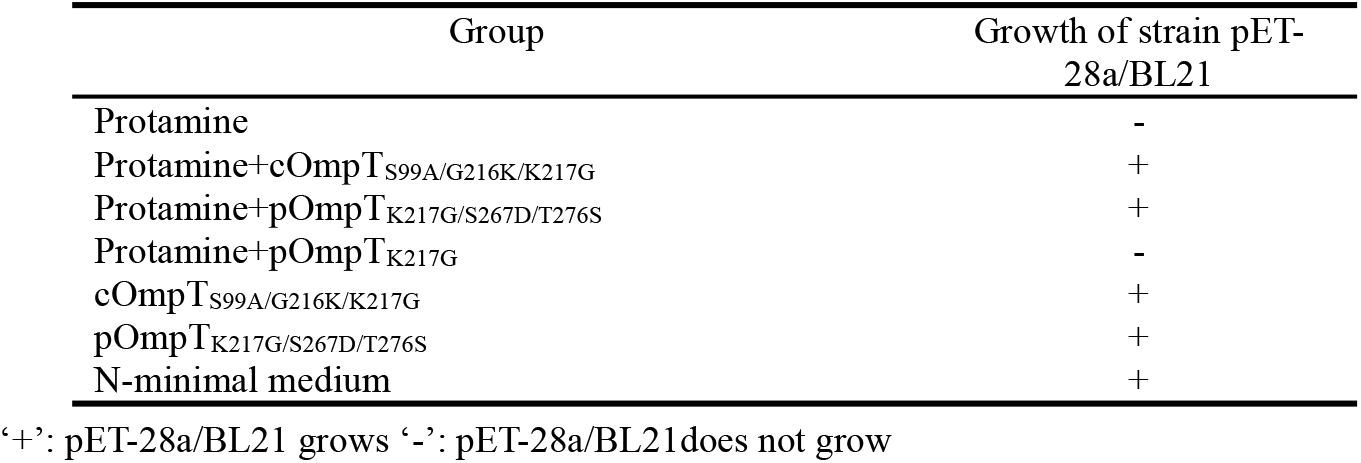
Identification of the activity and integrity of purified OmpT.

**Figure 3.**
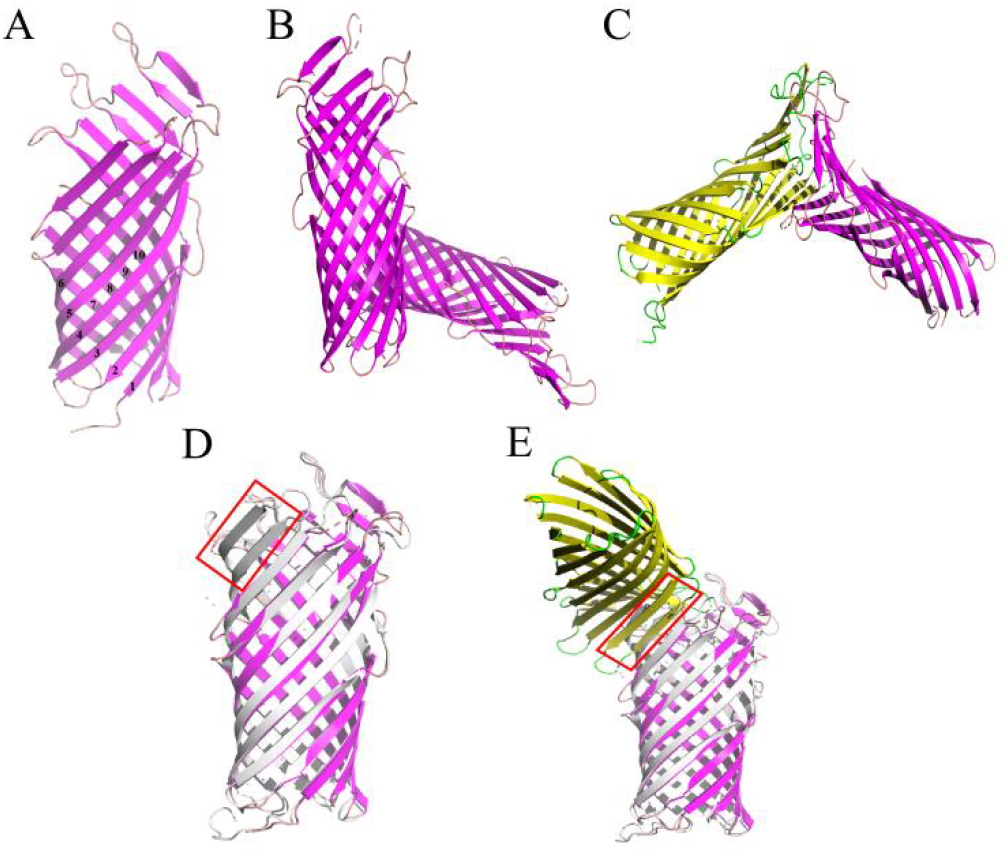
Crystal structure of pOmpT and superimposition with cOmpT. A One monomer of the asymmetric unit of pOmpT (pink). The arabic numbers represent the beta strands. B Two monomers of the asymmetric unit of pOmpT. C The monomer (maganeta) interacted with its symmetric molecules (yellow). D, E Superimposition of pOmpT (maganeta) with cOmpT (gray) (PDB No. 1I78). Yellow colored monomer in E represents the symmetric molecule interacting with pOmpT. Red box represents the missing region of pOmpT (residues 259-281).

### The difference of residues at positions 267 and 276 discriminates protamine cleavage by cOmpT and pOmpT

To test whether the structure missing region (in the loop 5) is important for the inactivation of protamine, we constructed mutants of pOmpT and cOmpT by exchanging the loop 5 of pOmpT and the corresponding region of cOmpT with each other. We evaluated the sensitivity of these constructs to protamine. Results indicated that the resistance to protamine of cOmpT construct with the fragment of pOmpT was dramatically decreased (Fig 2B). On the contrary, the resistance of pOmpT construct with a replaced loop 5 of cOmpT was dramatically increased (Fig 2B). We also constructed a series of mutants by exchanging other loop regions between cOmpT and pOmpT, protamine resistance assay results showed that they have no obvious effects. All above results indicated that the missing region was responsible for the difference of protamine resistance between cOmpT and pOmpT. Although the sequence identity between pOmpT and cOmpT reaches 78%, the sequence identity of the loop 5 is only 77.35%. In order to further identify the key residues of cOmpT on the loop 5 for the contribution in resisting protamine, the single/multiple residues of the loop 5 were exchanged between cOmpT and pOmpT. Protamine resistance assay showed that when the Ser^276^ of cOmpT was replaced with the Thr^276^ of pOmpT, the resistance to protamine was significantly reduced (*P*<0.05) (Fig 2C). When Asp^267^ of cOmpT is replaced with the Ser^267^ of pOmpT, or both Asp^267^ and Ser^276^ of cOmpT are replaced with Ser^267^ and Thr^276^ of pOmpT, the protamine resistance was extremely decreased (*P*<0.001) (Fig 2C). Compared to pOmpT, pOmpT_S267D_ significantly improved growth ability under resistance to protamine (*P*<0.05), and pOmpT_S267D/T276S_ significantly improved resistance to protamine (*P*<0.01) (Fig 2D). Growth kinetics of bacteria resistance to protamine further confirm the contribution of the residues 267 and 276 of cOmpT to the resistance to protamine in APEC E058 (Fig 2E and F).

We also analyzed protamine cleavage efficiency by *E. coli* with expressed cOmpT, pOmpT and their various mutants. The cleavage efficiency of protamine by single-mutant D267S and double-mutant D267S/S276T of cOmpT is significantly decreased in comparison to cOmpT (*P*<0.001) (Fig 4A and C). In contrast, the cleavage efficiency of protamine by single-mutant T276S and double-mutant S267D/T276S of pOmpT increased significantly (*P*<0.01) (Fig 4B and D). Notably the cleavage efficiency of single-mutant S267D of pOmpT is even higher than others (*P*<0.001) (Fig 4B and D). Above results indicated that the loop 5 of cOmpT is the key loop that contributes to different resistance to protamine between cOmpT and pOmpT and the key residues are residues 267 and 276.

**Figure 4.**
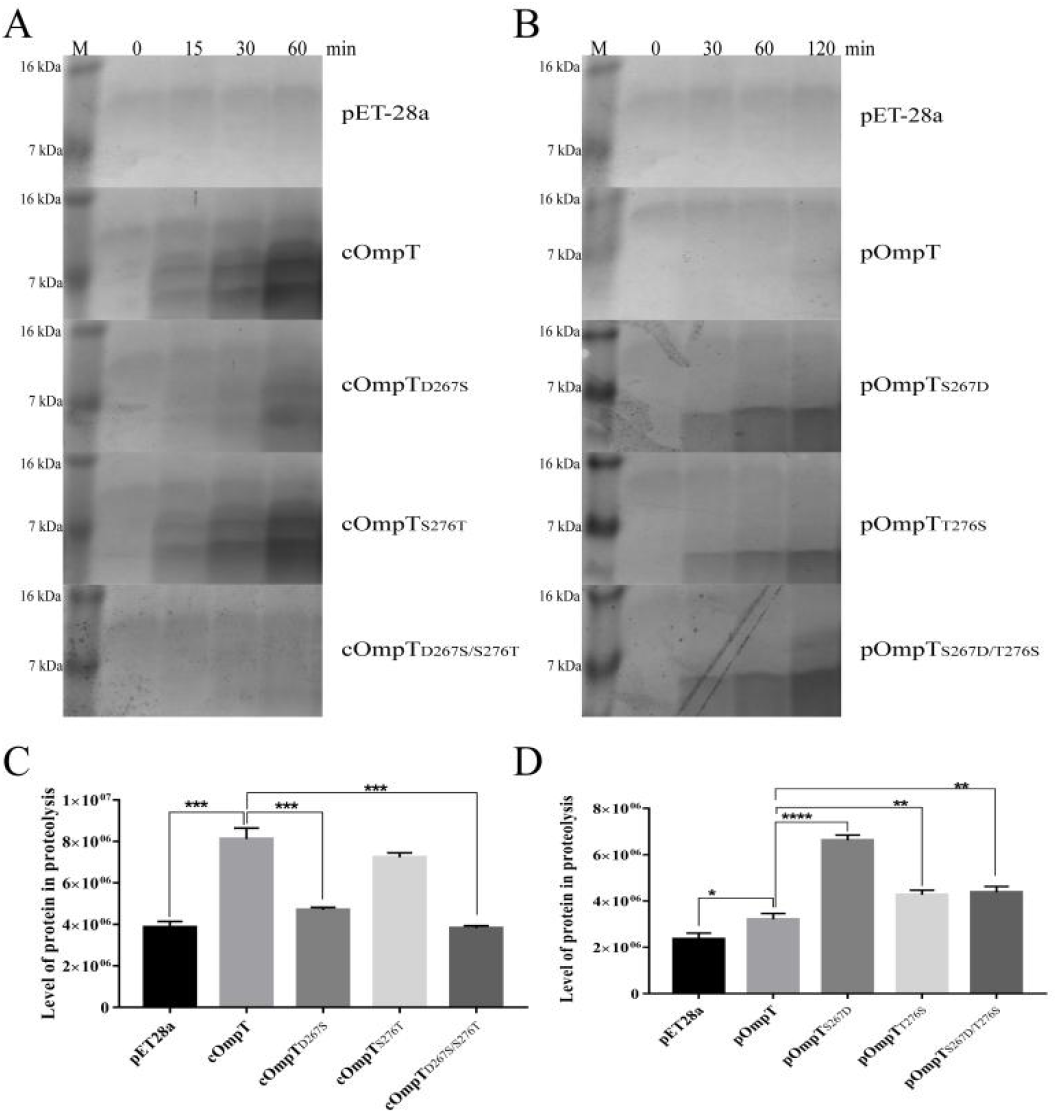
Cleavage of protamine by cOmpT, pOmpT and their site-directed mutants. A Protamine was incubated with *E. coli* BL21 expressing cOmpT (loops 1-5 of cOmpT) and its site-directed mutants (D267S and/or S276T) for the indicated times; B Protamine was incubated with *E. coli* BL21 expressing pOmpT(loops 1-5 of pOmpT) and its site-directed mutants(S267D and/or T276S) for the indicated times; C, D Quantitative analysis of the protamine products resolved by 16.5% Tris-Tricine SDS-PAGE and visualized with coomassie blue staining in a and b respectively. Data information: bars (C, D) represent mean values with standard error of the mean(n=3). Statistical significance was determinated using the *t*-test. Differences with *p*-values < 0.05 were considered as statistically significant. *: *p* < 0.05; **: *p* < 0.01; ***: *p* < 0.001; ****: *p* < 0.0001.

### Residues 267 and 276 of cOmpT and pOmpT are involved in substrate affinity

OmpT was reported to specifically cleave substrates at the dibasic motifs using synthetic peptide as substrate. We then checked whether cOmpT and pOmpT have same characteristics and whether the difference at residues 267 and 276 between cOmpT and pOmpT have effect on their activity to synthetic substrate (Dekker *et al*, 2001; McCarter *et al*, 2004). Digestion results showed that both pOmpT and cOmpT could cleave the synthetic substrate (Fig 5), albeit the cleavage efficiency of cOmpT is still higher than pOmpT, this result is different from the cleavage activity to substrate protamine which could only be digested by cOmpT. Single and double replacement mutation of cOmpT (S276T and D267S/S276T) dramatically decreased the activity of cOmpT (*P*<0.01), in contrast the replacement mutation of pOmpT significantly increased the cleavage efficiency in comparison to pOmpT (*P*<0.001) (Fig 5A). Interestingly, the highest cleavage efficiency protein is pOmpT_S276T_ emphasized the importance of this residue. These results further confirmed that residues 267 and 276 also contribute to the cleavage difference to synthetic substrate between cOmpT and pOmpT.

**Figure 5.**
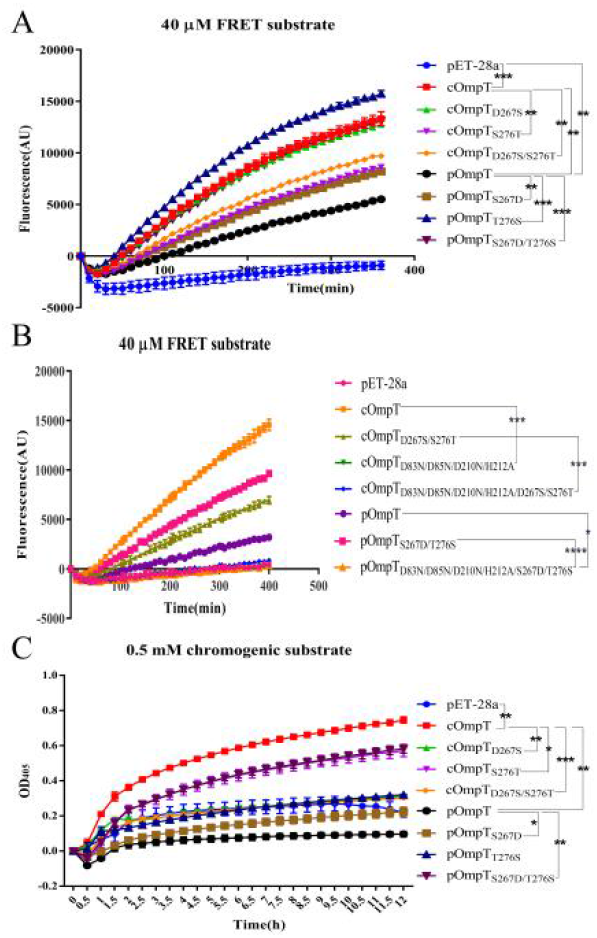
Determination of enzyme activity of cOmpT, pOmpT and its site-directed mutants. A, B Enzyme activity of cOmpT, pOmpT and their variants expressed in *E. coli* BL21 by utilizing a FRET substrate (2Abz-SLGRKIQI-K(Dnp)-NH2) C Enzyme activity of cOmpT, pOmpT and their variants expressed in *E. coli* BL21 by utilizing a chromogenic substrate (IAA-Arg-Arg-pNA). Data information: lines (A-C) represent mean values with standard error of the mean (n=3 reactions for each genotype). Statistical significance was determinated using the two-way ANOVA. Differences with *p*-values < 0.05 were considered as statistically significant. *: *p* < 0.05; **: *p* < 0.01; ***: *p* < 0.001; ****: *p* < 0.0001.

It was reported that cOmpT could be inhibited by serine protease inhibitor(Brannon *et al*, 2015). We test whether serine protease inhibitors inhibit pOmpT as cOmpT, results showed that pOmpT was also inhibited by both leupeptin and aprotinin (Fig 6G, K, H and L), and the mutants of residues 267 and 276 of c/pOmpT were also inhibited by the inhibitors (Fig 6), indicating that the difference between pOmpT and cOmpT does not account for the inhibition by the serine protease inhibitors. Residues (Asp^83^, Asp^85^, Asp^210^ and His^212^) were proposed as the catalytic residues for cOmpT (Kramer *et al*, 2001; Vandeputte-Rutten *et al*, 2001), the mutation of these residues of cOmpT, pOmpT and their mutants resulted in the loss of activity to synthetic substrate under the condition with or without the inhibitors suggested that these residues are also essential for the activity of pOmpT (Fig 5B; Fig 6C, D, G, H, K and L). These results also suggest that residues 267 and 276 may not be the catalytic residues for cOmpT and pOmpT. To further determinate the role of residues 267 and 276, we determined the *K*_m_ values of cOmpT, pOmpT and their mutants of residues 267 and 276. The *K*_m_ values of cOmpT mutants are all higher than the wild type cOmpT (1.56-4.44 times) (Table 2), indicating that the substrate binding affinity of these mutants is decreased. In contrast, the *K*_m_ value of pOmpT is higher than its mutants (2.37-4.24 times) (Table 2). The above results suggest that residues 267 and 276 are involved in the substrate binding rather than catalysis. We also determined the *K*_cat_/*K*_m_ of c/pOmpT and their 267 and 276 mutants, it is obvious that the catalytic efficiency of cOmpT is much higher than pOmpT, the mutation of residues 267 and 276 improved the catalytic efficiency of pOmpT, in contrast, the corresponding mutation of cOmpT decreased the catalytic efficiency dramatically (Table 2), the different catalytic efficiency between c/pOmpT and their mutants indicated that the binding affinity change caused by residues 267 and 276 further affects the catalytic efficiency.

**Table 2.**
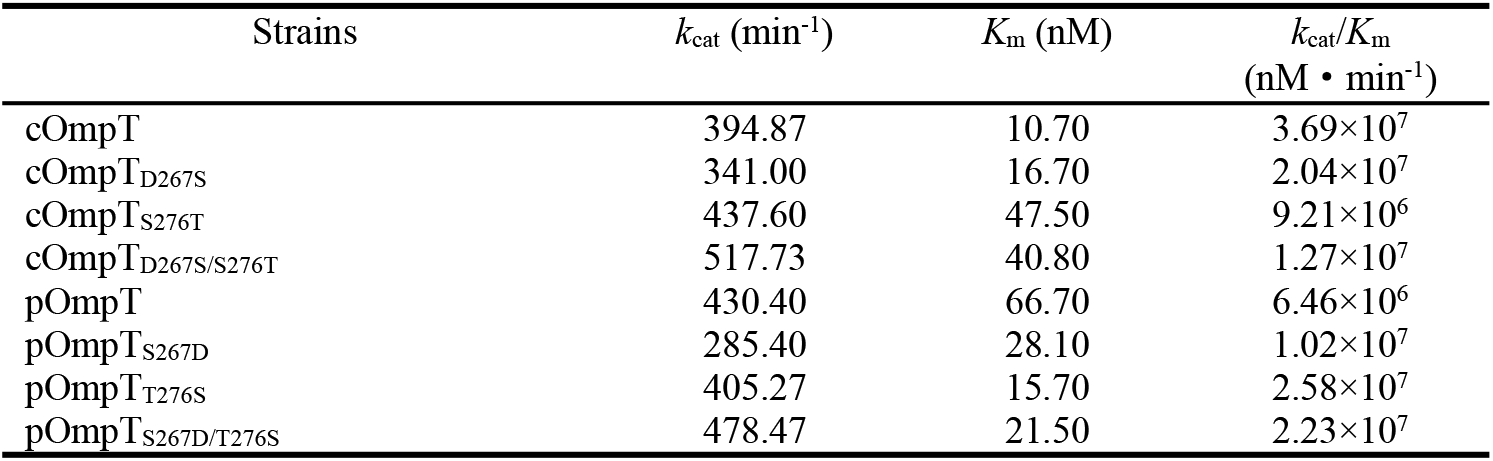
Kinetic parameters of (c/p)OmpT and their site-directed mutants expressed in *E. coli* BL21 using synthetic fluorimetric peptide as substrate.

**Figure 6.**
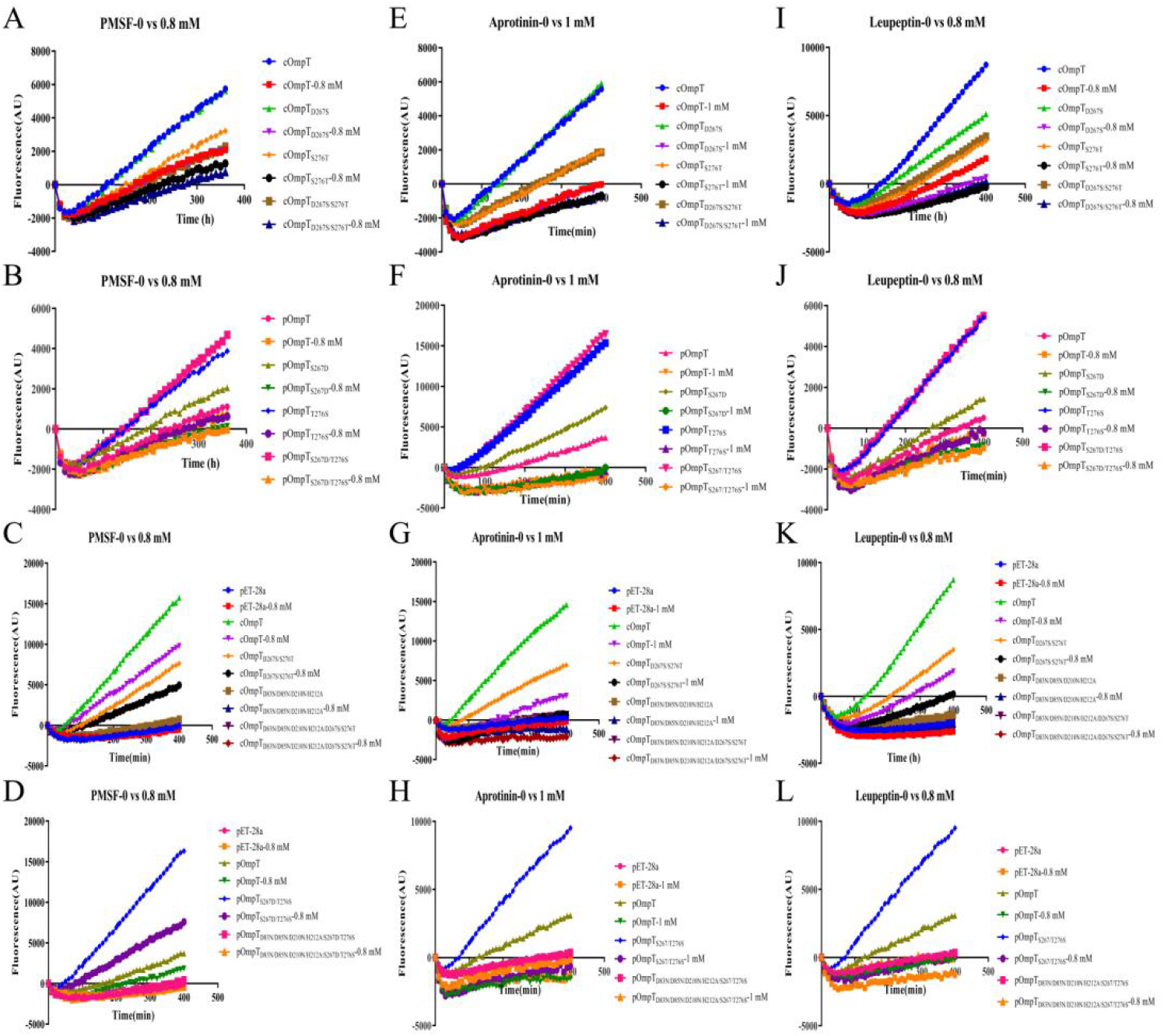
Inhibition of enzyme activity of cOmpT, pOmpT and its site-directed mutants expressed in *E. coli* BL21. A-D FRET assays were performed with cOmpT, pOmpT and its site-directed mutants in PBS or in the presence of PMSF (0.8 mM). E-H FRET assays were performed with cOmpT, pOmpT and its site-directed mutants in PBS or in the presence of aprotinin (1 mM). I-L FRET assays were performed with cOmpT, pOmpT and its site-directed mutants in PBS or in the presence of leupeptin (0.8 mM).

### Changes of residues at positions 267 and 276 of pOmpT can also change its ability to cleave RNase 7

ArlC in UPEC, which shares 100% identity with pOmpT, can promote bacterial resistance to the host by cleaving large molecule AMP human RNase 7 (Desloges *et al*, 2019). As expected, RNase 7 was cleaved when incubated with the strain expressed pOmpT as compared with the strains without c/pOmpT (such as strains E058Δc*ompT*Δp*ompT* and empty vector), these strains did not cleave RNase 7 with a clear RNase 7 protein band left (∼17.7 kDa), indicating pOmpT cleaved RNase7 as ArlC (Fig 7A and B). Interestingly, here RNase 7 was also completely cleaved by cOmpT which was shown with no activity on RNase 7 in UPEC isolate cystitis 6 (Desloges *et al*, 2019). Synthetic substrate digestion showed that residues 267 and 276 of c/pOmpT are involved in the substrate recognition (Fig 7 and Table 2), so we also checked the influence on the proteolytic activity of c/pOmpT on RNase 7 by the change of residues 267 and 276. The result showed that the replacement of residues 267 and 276 of cOmpT with that of pOmpT dramatically decreased the activity of cOmpT on RNase 7, a clear band of intact RNase 7 was observed compared with cOmpT (Fig 7A). However, the replacement of these two residues of pOmpT with that of cOmpT did not have obvious effect on the activity of pOmpT (Fig 7A). One possible reason for this result may be that pOmpT is expressed at relative higher level due to its high copies coded by plasmid. To further check whether the replacement of residue 267 and 276 of pOmpT with that of cOmpT, we expressed cOmpT, pOmpT and their variant mutants in *E. coli* BL21 which does not have OmpT natively. Cleavage efficiency of cOmpT is clearly higher than pOmpT (Fig 7B). The replacement of residues 267 and 276 of cOmpT with that of pOmpT decreased the activity of cOmpT (Fig 7B). In contrast, the replacement of residues 267 and 276 of pOmpT with that of cOmpT increased the activity of pOmpT (Fig 7B). Mutation of residues 267 and 276 further emphasized the importance of residues 267 and 276 on the activity of c/pOmpT. An interesting observation is that the replacement of whole loop 5 of cOmpT with that of pOmpT increased the digestion activity to RNase 7 compared with the decreased activity caused by the replacement of residues 267 and 276 (Fig 7B).

**Figure 7.**
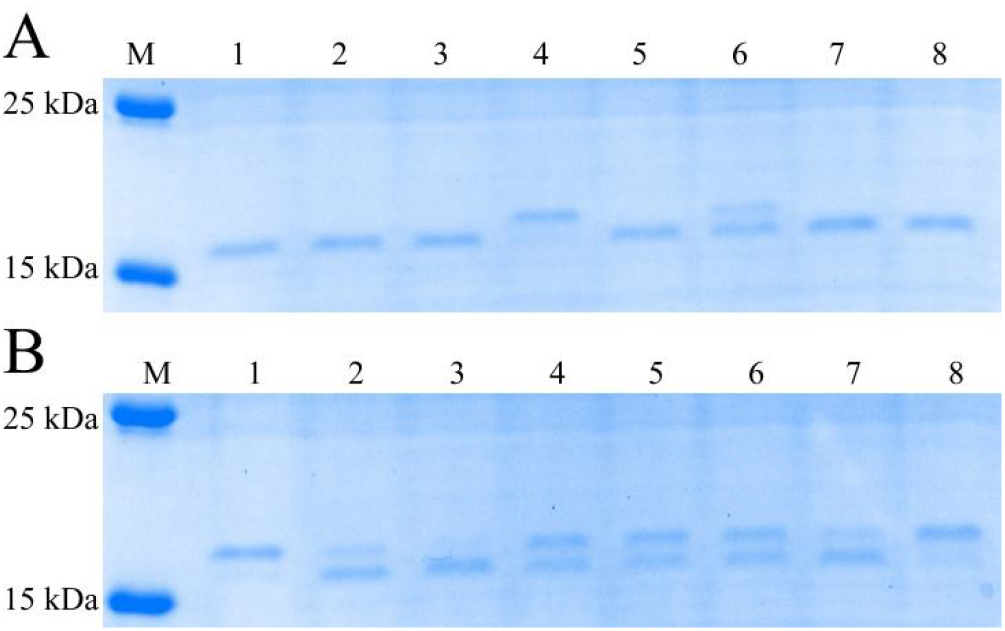
Cleavage of RNase 7 by cOmpT, pOmpT and their mutants. A RNase 7 was incubated with APEC E058 expressing cOmpT, pOmpT or their mutants for 15 min; Lane 1: E058; Lane 2: E058Δc*ompT*; Lane 3: E058Δp*ompT*; Lane 4: E058Δc*ompT*Δp*ompT*; Lane 5: E058c*ompT*_c1-4p5_Δp*ompT*; Lane 6: E058c*ompT*_D267S/S276T_Δp*ompT*; Lane 7: E058p*ompT*_p1-4c5_Δc*ompT*; Lane 8: E058p*ompT*_S267D/T276S_Δc*ompT*; B RNase 7 was incubated with *E. coli* BL21 expressing cOmpT, pOmpT or their variants for 15 min; Lane 1: empty vector pET-28a; Lane 2: cOmpT; Lane 3: c1-4p5; Lane 4: cOmpT_D267S/S276T_; Lane 5: pOmpT; Lane 6: p1-4c5; Lane 7: pOmpT_S267D/T276S_; Lane 8: Negative Control; 180 kDa prestained protein marker (Vazyme, China) was used as the protein molecular size standard (lane M). Detailed information of bacterial strains in Expanded View Table EV1.

## Discussion

Omptins are key virulence factors involved in several Gram-negative bacterial pathogens(Sodeinde, Subrahmanyam et al. 1992, Lathem, Price et al. 2007, Caulfield, Walker et al. 2014, Mcphee, Small et al. 2014). Numerous studies suggested that OmpT-like proteins have been found in the intestinal or extra-intestinal pathogenic *E. coli* that infect humans and animals, and they enhance bacterial virulence by participating in the cleavage of host AMPs (Thomassin *et al*, 2012; Brannon *et al*, 2013; Desloges *et al*, 2019; Liu *et al*, 2019). Their contribution to *E. coli* virulence mainly depends on their own proteolytic activity and substrate specificity (Hritonenko & Stathopoulos, 2007). It has been confirmed that Omptins in *E. coli* mainly include OmpT encoded on nucleoids and OmpP/ArlC encoded on episomal plasmids (Hwang *et al*, 2007; Mcphee *et al*, 2014; Desloges *et al*, 2019). Most pathogenic *E. coli* (eg, NMEC, EPEC, EHEC, UPEC) only have OmpT encoded on the nucleoid. However, here we found that APEC E058 strain (O2 serotype) belonging to the ExPECs harbored the p*ompT* located on the *ColV* plasmid in addition to the c*ompT* located on the nucleoid. Surprisingly, although the amino acid sequence identity of these two homologs is only 78%, both of them are expressed on the outer membrane of APEC E058. In addition, the characterized active sites (including catalytic residues) and LPS-binding sites of cOmpT are conserved in pOmpT except for the two LPS-binding sites. Sequence alignment revealed that both APEC cOmpT and UPEC OmpT (from UPEC CFT073 and UPEC cystitis isolate 6) were derived from the same ancestor, while pOmpT in APEC and ArlC in UPEC cystitis isolate 6 were homologs based on this study. OmpT is able to cleave protamine (Stumpe *et al*, 1998), which was confirmed in APEC E058. ArlC was first detected in the pathogenicity island on the plasmid of the adherent-invasive *Escherichia coli* (AIEC) NRG 857c strain and involved in the resistance to the human host defense peptides HD5, HBD2 and LL-37 (Mcphee *et al*, 2014). Later, ArlC was also found in UPEC cystitis isolate 6 with the ability to cleave human RNase 7 (Desloges *et al*, 2019). However, whether pOmpT or ArlC can resist protamine remains unclear. Here, the findings that pOmpT unable to cleave protamine fills the gap.

OmpT-like proteases may differ in substrate specificity, suggesting significant structural differences in the proteolytic active site enriched grooves between cOmpT and pOmpT despite their high amino acid sequence identity (Brannon *et al*, 2015). Therefore, we try to explore the crystal structure of pOmpT. Previously, the only resolved structure of *E. coli* OmpT-like subfamily protease was cOmpT encoded on the nucleoid, which displayed a 10-chain antiparallel β-barrel, and its extracellular loop protrudes above the lipid bilayer (Vandeputte-Rutten *et al*, 2001). Putative binding sites for LPS and the active sites (including the catalytic sites) are identified in the groove (substrate-binding pocket) at the extracellular top of the vase-shaped β-barrel (Vandeputte-Rutten *et al*, 2001). The crystal structure of pOmpT we resolved here is largely similar to that of cOmpT, also presents a typical OmpT fold, a vase-shaped antiparallel β-barrel. However, data from residues 259 to 281 located in the extracellular loop of pOmpT were missing due to the conformational flexibility, leading to a significant difference of the substrate-binding pocket of pOmpT, ultimately, resulting in a loss of protamine cleavage by pOmpT. To trace the resistance of protamine to pOmpT, the effect of the structurally missing region of pOmpT (residues 259 to 281) on pOmpT proteolytic activity and substrate specificity was focused. Previous study revealed that the proteolytic activity and substrate specificity of omptins depend on the residues in its loop 1-5 (L1-L5), and mutation of these residues alters its proteolytic activity (Kukkonen *et al*, 2001). The crystal structure analysis of cOmpT confirms that the active sites are actually distributed in the substrate-binding pocket region at the top of the extracellular loop (Vandeputte-Rutten *et al*, 2001). The kinetics of protamine cleavage by loop-swap mutants suggested that loop 5 (L5) of cOmpT contributes to the activity of protamine cleavage. Further, Asp^267^ and Ser^276^ in L5 of cOmpT are identified as the key residues to cleave protamine. Since active sites and putative LPS-binding sites of cOmpT have been characterized, neither 267^th^ nor 276^th^ residue has been involved in these active sites. Therefore, our findings are particularly important to supplement the critical information.

The *E. coli* outer membrane protease OmpT was previously classified as a serine protease, since its Ser^99^ and His^212^ are confirmed as typical active sites (Kramer *et al*, 2000). Later, OmpT was also identified as an aspartic protease (Vandeputte-Rutten *et al*, 2001), and a novel proteolytic mechanism that involves a His^212^-Asp^210^ dyad and an Asp^83^-Asp^85^ pair that activate a putative nucleophilic water molecule has been proposed based on the crystal structure of *E. coli* OmpT (Vandeputte-Rutten *et al*, 2001). The active sites are fully conserved in the omptin family (Vandeputte-Rutten *et al*, 2001). Here we focus whether pOmpT experiences the same proteolytic mechanism as cOmpT and the effect of residues 267 and 276 on proteolytic activity. Sequence alignment analysis of cOmpT and pOmpT revealed that their catalytic residues are conserved between the two homologous. We confirmed that once these four catalytic sites (Asp^83^, Asp^85^, Asp^210^ and His^212^) were mutated, the proteolytic activity of the mutants was completely abolished, neither 267 nor 276 residue interchange between cOmpT and pOmpT was able to eliminate the ability to cleave substrates, implying that these two residues were not catalytic sites of OmpT. Earlier study supported the hypothesis that OmpT is a member of the serine protease family (Kramer *et al*, 2000), since the proteolytic activity of OmpT can be significantly inhibited by serine protease inhibitors (such as diisopropylfluorophosphate (DFP), phenylmethanesulfonyl fluoride (PMSF)) (Sugimura & Nishihara, 1988). In line with the observation in cOmpT, the proteolytic activity of wild-type pOmpT and its variant S267D/T276S were significantly inhibited by serine protease inhibitors (Aprotinin, PMSF and Leupeptin), suggesting a catalytic mechanism shared by pOmpT and cOmpT. We then analyzed the enzymatic activities of wild-type cOmpT, pOmpT and their substitutions at residues 267 and 276 using synthetic substrates. The proteolytic activity of pOmpT was found to be significantly lower than that of cOmpT, corresponding to the inability of pOmpT to cleave protamine. Substitutions with either S276T or D267S/S276T significantly impairs the proteolytic activity of wild type cOmpT, on the contrary, either S267D, T276S or S267D/T276S replacement significantly enhances the proteolytic activity of wild type pOmpT. These results fully demonstrated that residues 267 and 276 of cOmpT and pOmpT are active sites contributed to the proteolytic activity of both cOmpT and pOmpT, leading to different substrate cleavage specificities recognized by these two omptins. Despite the active sites of Omptins are fully conserved (Hritonenko & Stathopoulos, 2007), neither residue 267 nor 276 located at L5 of both cOmpT and pOmpT has been proposed as active sites in our knowledge. To further clarify the role of residues 267 and 276, proteolytic kinetics of cOmpT, pOmpT and their mutants involved in residues 267 and 276 were carried out using synthetic fluorescent peptides. Compared to the wild-type cOmpT, cOmpT mutants displayed essentially undiminished values for the catalytic constant (*k*_cat_), but the *K*_m_ values of the mutants were increased by approximately 56% (Asp^267^-to-Ser substitution, (16.7-10.7)/10.7), 344% (Ser^276^-to-Thr substitution, (47.5-10.7)/10.7) and 281% (both Asp^267^ and Ser^276^ substitutions, (40.8-10.7)/10.7), respectively. In contrast, compared to the wild-type pOmpT, the *K*_m_ value of pOmpT mutants were reduced by approximately 58% (Ser^267^-to-Asp substitution, (66.7-28.1)/66.7), 76% (Thr^276^-to-Ser substitution, (66.7-15.7)/66.7) and 68% (both Ser^267^ and Thr^276^ substitutions, (66.7-21.5)/66.7), respectively. These indicated that residues 267 and 276 of cOmpT and pOmpT are involved in the substrate binding rather than in catalysis. In accordance with Tyr^248^ in carboxypeptidase A, Tyr^248^ did not contribute to catalysis, instead, involved in substrate binding, since the Tyr^248^-to-Phe substitution kept the same *k*_cat_ value while its *K*_m_ value was increased 6 times compared to the wild type (Gardell *et al*, 1985). Therefore, we inferred that residues 267 and 276 of cOmpT and pOmpT functioned as substrate binding sites, involving in anchor and ultimate promotion of the substrate cleavage by the catalytic center.

The presence of two OmpT-like proteases in UPEC clinical isolates may confer a fitness advantage by expanding the range of target substrates during urinary tract infections, among them, (c)OmpT can cleave LL-37, while ArlC can specifically cleave RNase 7 (Desloges *et al*, 2019). However, we found that both cOmpT and pOmpT of APEC E058 strain origin were able to cleave RNase 7. More importantly, in accordance with enzymatic activities determined using synthetic substrates, substitutions of residues 267 and 276 between cOmpT and pOmpT similarly affected their ability to cleave RNase 7. It is interested that pOmpT of APEC E058 strain dose not cleave protamine while does cleave RNase 7, although both of them belonged to human AMPs. OmpT protease has a narrow cleavage specificity, showing preferential cleave substrates at dibasic motifs (Dekker *et al*, 2001; McCarter *et al*, 2004). This specificity of OmpT is determined by conserved Glu^27^ and Asp^208^ at the bottom of the deep S1 pocket and Asp^97^ in the shallower S1 pocket (Vandeputte-Rutten *et al*, 2001). However, pOmpT and cOmpT are highly conserved at the above three residues, suggesting that residues 267 and 276 of cOmpT play a key role in substrate binding through interaction with arginine residues in protamine. Based on OmpT-Pla chimeric protein, substrate specificity differs between different omptins through sequence variability in the outer loop (Kukkonen *et al*, 2001). Here the structural flexibility of the missing region on the 5-loop of pOmpT and the difference in target substrates (e.g., protamine and RNase 7) recognition by residues 267 and 276 reveal the substrate binding specificity of pOmpT. Therefore, we speculate that the specificity of the structure of pOmpT prevents it from interacting with arginine residues in protamine to initiate the proteolytic process, instead, cleave RNase 7 after effectively binding other amino acids in RNase 7.

Currently, there are many bacterial infections without effective treatment or prevention due to the more and more antibiotic resistance. Omptins might offer new potential targets for drug and vaccine development, and inhibition of these omptins is critical in preventing septisemic bacterial infections (Brannon *et al*, 2015; Hritonenko & Stathopoulos, 2007). Here we characterized the structure and physiology of pOmpT, a newly discovered OmpT-like subfamily protease in pathogenic *E. coli* caused intestinal and extraintestinal infections in both humans and poultry, and revealed its structural basis and molecular mechanism involved in human AMPs cleavage. Residues 267 and 276 were firstly characterized as substrate binding sites of omptins, and playing a critical role in efficiency of AMPs cleavage by cOmpT and pOmpT. Importantly, both cOmpT and pOmpT of APEC focused on this study are homologs of human ExPEC, especially, cOmpT and pOmpT of avian origin confers APEC with ability to cleave human AMPs, supporting that APEC are potential human pathogens (Johnson *et al*, 2008). Our findings provide new clues for the development of antibacterial drugs target inhibition of omptins activity.

## Materials and Methods

### Bacterial strains, plasmids and growth conditions

Bacterial strains and plasmids used in this study are listed in Expanded View Table EV1. Oligonucleotide primers are listed in Expanded View Table EV2. APEC strain E058 was isolated from a chicken with the typical clinical symptoms of colibacillosis in China (Gao *et al*, 1999). The other details of strains are as follows. The strains were grown in Luria-Bertani (LB) broth or on LB agar plates or in N-minimal medium (Nelson & Kennedy, 1971) adjusted to pH 7.5 and supplemented with 0.2% glucose and 1 mM MgCl_2_. Antibiotics like 50 μg/mL kanamycin, 50 μg/mL spectinomycin, 30 μg/mL chloramphenicol or 50 μg/mL tetracycline were used for selection wherever required. All the cultures were grown at 37°C or 30°C under aerobic conditions.

### Generation of mutants, revertants and recombinant expression bacteria

The single/double gene deletion of c*ompT* and p*ompT* genes of APEC E058 strain was constructed using lambda red recombinase system (Liu *et al*, 2019). At the same time, both the native c*ompT* and p*ompT* genes, along with their native putative promoters, were amplified and cloned into plasmid pACYC184. After PCR and DNA sequencing, the correct plasmids p184-cc*ompT* and p184-pp*ompT*, were correspondingly transformed into the c*ompT*/p*ompT* single gene deletion strains to generate the complementation strains by electroporation.

The c*ompT* and p*ompT* genes without signal peptide and with His tag, and with site-directed mutations were amplified by the overlap PCR, cloned into the expression plasmid pET-30a after digestion, and finally transferred to bacteria BL21. In order to reduce the interference of the outer membrane proteins other than the cOmpT/pOmpT protein in the E058 strain, the amplified c*ompT* and p*ompT* genes of the whole open reading frame (ORF) were cloned into the expression plasmid pET-28a, and the correct plasmids were transferred into BL21 after PCR and DNA sequencing. In addition, the amino acid sequences of cOmpT and pOmpT proteins are divided into 5 loops according to the topological structure of cOmpT protein. Combined with the fusion PCR technology, we used the same gene cloning method as above to construct a series of recombinant expression bacteria with corresponding interchanges of 5 loops and site-directed mutations in the loop 5.

In order to further determine the importance of the loop 5 and the amino acid at positions 83/85/210/212/267/276 of cOmpT/pOmpT in the wild-type strain E058, the mutual interference between cOmpT and pOmpT must be excluded. Therefore, we first realize the insertion of the chimera or site-directed mutant DNA fragments at the gene deletion and then realize the deletion of another gene on the basis of the c*ompT*/p*ompT* single gene deletion strain using a CRISPR-Cas9 system-based continual genome editing strategy (Jiang *et al*, 2015). The guide sequences (N20 sequence) were used to construct pTarget series plasmids, which is respectively target to the FRT sequence of c*ompT*/p*ompT* single gene mutant strain, c*ompT* and p*ompT* gene sequences. The donor DNA were amplified correspondingly using the genome DNA of the c*ompT*/p*ompT* single-gene deletion strain and above correct pET series plasmids with loop 5 interchange and amino acid site mutation/interchange of the c*ompT*/p*ompT* gene as templates.

The DNA template of RNase 7 was derived from the experimenter’s nasal swab sample. The *rnase 7* gene without signal peptide, and with His tag and site-directed mutations was amplified by the overlap PCR and cloned into the expression plasmid pDEST17 after digestion. Then, the correct plasmid pDEST17-RNase 7 was transformed into competent cells BL21(AI) (Koten *et al*, 2009; Wang *et al*, 2013).

### RNA isolation, reverse transcription (RT)-PCR, sequencing and alignment analysis

Total RNA was extracted from APEC strain E058 and reverse-transcribed into cDNA by using the PrimeScript RT reagent kit (TaKaRa, China) according to the manufacturer’s protocol. Primer sets for PCR amplification of the target gene c*ompT* and p*ompT* in cDNA samples are shown in Expanded View Table EV2. In parallel, PCRs were performed with strain E058 DNA as the positive controls and cDNA samples without activation of the reverse transcription (RT) as negative controls. The PCR products were resolved on 0.8% agarose gels. Then the fragments corresponding to the PCR-amplified gene c*ompT* and p*ompT* were extracted from the agarose gels using an Axygen DNA gel extraction kit (Corning, China), and were sent for sequencing verification. Finally, sequence alignment is performed among the amino acid coding sequence deduced from the sequenced APEC E058 c*ompT* and p*ompT* gene’s nucleic acid sequences, the amino acid coding sequence deduced from the *ompT* (GenBank: CP051263.1) gene’s nucleic acid sequence in the UPEC CFT073 and the nucleic acid sequence of the *ompT* (GenBank: CP041302.1) and *arlC* (GenBank: CP041301.1) genes in the cystitis 6 isolate published on NCBI.

### Outer membrane protein extraction and western blotting

Bacteria were cultured overnight in 100 mL LB broth medium. Outer membrane fractions were isolated as follows (Gao *et al*,, 1996): bacterial cells were centrifuged at 6,000 rpm for 10 min at 4°C and pellets were resuspended in 6 mL HEPES buffer (10 mM HEPES, pH 7.4) and sonicated. Samples were then centrifuged at 6,000 rpm for 10 min at 4°C. Supernatants were collected and supplemented with about 8 times the volume of sarcosyl buffer (2% sarcosyl) and incubated for 30 min at 4°C. Samples were then centrifuged for 1 h at 35,000 rpm, and the pellet containing outer membranes was resuspended in 1 mL buffer A (20 mM Tris-HCl pH 7.5 and 10% glycerol). Finally, the concentration of the extracted total outer membrane protein was determined with the BCA protein concentration determination kit (Beyotime, China). Outer membrane samples were combined 1:3 with 4× protein loading buffer (Vazyme, China) and boiled for 10 min. Outer membrane fractions were resolved on a 12% SDS-PAGE gel and transferred to a polyvinylidene fluoride membrane. Membranes were blocked overnight in PBST buffer (10 mM phosphate-buffered saline [PBS, pH 7.4], 0.05% Tween-20) supplemented with 5% skim milk at 4°C, and OmpT was detected using the monoclonal anti-OmpT antibody. Membranes were washed extensively with PBST and incubated for 1 h with a goat anti-mouse secondary antibody conjugated with HRP. Membranes were washed and developed using chemiluminescent HRP substrate.

### Protein expression and purification of cOmpT, pOmpT and RNase 7

The expression and purification of cOmpT and pOmpT were carried out on the basis of predecessors (Sugimura & Nishihara, 1988; Vandeputte-Rutten *et al*, 2001). The BL21(DE3) strain harboring the expression plasmid pET-cOmpT_S99A/G216K/K217G_, pET-pOmpT_K217G_ and pET-pOmpT_K217G/S267D/T276S_ was inoculated in LB broth medium at 37°C until the optical density at 600 nm (OD_600_) reached approximately 0.6–0.8. Then, the cells were induced with isopropyl β-d-1-thiogalactopyranoside (IPTG) at a final concentration of 0.4 mM and grown at 37°C for approximately 5 h. The induced cells were harvested by centrifugation at 5000 rpm for 10 min at 4°C, and the pellet was resuspended in the lysis buffer (20 mM Tris-HCl, 2%Triton X-100, 1 mM EDTA, pH 8.4). Then, PMSF was added to a final concentration of 1 mM before sonication, the resuspended cells were lysed by ultrasonic disruption (power 5 W, ultrasound for 3 s, stop for 3 s). The lysed precipitate was collected by centrifugation at 12000 rpm for 15 minutes, and washed twice with washing buffer (20 mM Tris-HCl and 1 mM EDTA, pH 8.4), and finally resuspended with washing buffer at a ratio of 1 mL washing buffer per precipitate of 1 L cell culture. The collected precipitate was resuspended with 6 mL of denaturation buffer (50 mM glycine and 1 M urea) at 4°C with slow shaking for 30 min. Then pre-chilled 240 mL of refolding buffer (10 mM PBS and 1% N-Dodecyl-N, N-dimethyl-3-ammonio-1-propanesulfonate) was quickly added in resuspension and slowly shaked at 4°C to refold for 12 h. Refolding buffers with different pH values were formulated due to the different isoelectric points of cOmpT and pOmpT. cOmpT_S99A/G216K/K217G_ was renatured with the refolding buffer with a pH of 5.3, while the pH of the refolding buffer used for pOmpT_K217G_ and pOmpT_K217G/S267D/T276S_ was 7.3. The refolded supernatants were collected by centrifugation at 12000 rpm for 15 minutes and place it in an ice-water mixture for later use. The protein purification procedures were carried out at 4°C as follows. Refolded native cOmpT_S99A/G216K/K217G_, pOmpT_K217G_ and pOmpT_K217G/S267D/T276S_ were loaded onto an 80 ml Fast Flow S-Sepharose column (GE Healthcare, USA) pre-equilibrated with buffer A (1% N-Dodecyl-N, N-dimethyl-3-ammonio-1-propanesulfonate (Sigma, USA) and 20 mM sodium acetate). The column was washed with buffer A and proteins were eluted with a linear gradient of NaCl to1 M in buffer A. The native protein was concentrated to less than 1 mL in buffer B (1% OG, 20 mM Tris-HCl, 150 mM NaCl, and 5% glycerol) by Amicon Ultra-50 filter (Millipore, USA). Then the concentrated target protein was loaded onto a Superdex 200 column pre-equilibrated with buffer B, and the protein was eluted with the same buffer. After concentrate and dialysis, samples used for crystallization consisted of 30 mg/mL pOmpT in buffer C (1% OG, 2.5 mM potassium chloride and 5 mM sodium acetate). cOmpT_S99A/G216K/K217G_ was purified with buffers A, B and C with a pH of 4.0, while buffers A, B and C used for pOmpT_K217G_ and pOmpT_K217G/S267D/T276S_ had a pH of 6.0. The target protein was analyzed by sodium dodecyl sulfate-polyacrylamide gel electrophoresis (SDS-PAGE) and concentrated and stored at 4°C until use.

RNase 7 was expressed in *E. coli* BL21(AI) harboring the expression plasmid pDEST17-RNase 7. 2 L of cultures were induced with L-arabinose (final concentration 2%) in LB broth medium for 3 h. The cell pellet was collected by centrifugation and resuspended in 20 mL buffer D (20 mM Tris-Cl, pH 7.5). After cells lysed by sonication, the supernatant was collected by centrifugation and purified through a Ni-NTA column (eluent: 50 mM Na_2_HPO_4_, 0.3 M NaCl and 250 mM imidazole, pH 8.0) and a Superdex-200 column (buffer E: 20 mM Tris-HCl, 150 mM NaCl and 10% glycerol, pH 8.0). RNase 7 is stored in buffer E (20 mM Tris-HCl, 150 mM NaCl, 10% glycerol) at 4°C. The protein concentration determined by BCA is about 0.15 mg/mL.

### Identification of the activity and integrity of purified OmpT

100 μL of purified cOmpT_S99A/G216K/K217G_, pOmpT_K217G_ and pOmpT_K217G/S267D/T276S_ and protamine (final concentration 100 μg/mL) were cultured in N-minimal medium with shaking at 37°C for 2 h. Then, the pET-28a/BL21 strain was added to each of the above culture reactions, and the culture was continued for 12 h. The growth of bacteria in the culture reaction is used as an indicator to identify the activity and integrity of the purified protein.

### Protein crystallization of pOmpT

Crystallization screening was carried out at 4 and 20°C employing the sitting drop vapor diffusion method. Initial crystallization screening was carried out using the condition of Lucy Vandeputte-Rutten’s crystallization condition for cOmpT and 10 crystallization kits from Hampton Research (Index, MembFac, Crystal Screen, Crystal Screen 2, Crystal Screen Lite and Crystal Screen Cryo) and Microlytic (MSG1, MSG2, MSG3 and MSG4). Crystal screening was performed with protein concentrations of 18, 19, 20, 21 and 22 mg/mL. The crystallization drop was set up with 1 μL of protein solution and 1 μL of reservoir solution, and the droplet was equilibrated against 500 μL of reservoir solution. The initial screening produces rhombic crystals under multiple conditions: pOmpT 18-22 mg/mL, 28-31% MPD, 0.3 M sodium citrate (pH 5.3-5.8). The initial condition was further optimized by adjusting the concentration of MPD and the pH. The best diffraction crystal was obtained under the condition of protein concentration 20 mg/mL and 30% MPD (pH 5.5). The crystals grew to their maximum size after 6 months.

### Data collection, processing, and structure determination

Before data collection, the crystals were looped out and quickly dipped in liquid nitrogen and preserved in a liquid nitrogen tank. Data collection was performed at the Shanghai Synchrotron Radiation Center using beamlines BL18U1. Diffraction data were processed using the X-ray Detector Software (XDS) package (Kabsch, 2010) and scaled by SCALA (Evans, 2006). The structure of the pOmpT-K217G was determined by the molecular replacement method using the program Phaser (Mccoy *et al*, 2007) using the structure of cOmpT (Protein Data Bank with accession No.1I78) from *E. coli* (Vandeputte-Rutten *et al*, 2001) as the search model. Manual model correction was performed using Coot (Emsley & Cowtan, 2004). Refinement was performed using REFMAC5 (Murshudov *et al*, 1997) and Phenix (Adams *et al*, 2010).

### Growth kinetics of bacteria incubated with protamine

In order to study the growth kinetics of bacteria incubated with protamine, we chose N-minimal medium to cultivate bacteria in order to avoid the potential interference by some components of complex medium on the function of OmpT. Overnight bacterial cultures in N-minimal medium were prepared. The cultures were centrifugated and resuspended in 10 mL fresh N-minimal medium and normalized to an OD_600_ nm of 0.35 (a final concentration of 3.5 × 10^8^ CFU/ml). Protamine was then added to the bacterial suspension at a final concentration of 100 μg/mL. The mixture was shaken at 37°C and measured the value of OD_600_ nm every 1 or 2 h. The results were drawn into the growth curve of bacteria, and used GraphPard Prim 7 software for differential analysis.

### Proteolytic cleavage of AMPs

Bacterial cells were grown in N-minimal medium to an OD_600_ nm of 0.6-0.8. Then the culture was washed, pelleted by centrifugation, resuspended in PBS (pH 7.4) and normalized to a bacterial density of 3× 10^10^ CFU/mL. Aliquots of bacteria were combined 1:4 (v/v) with 2.5 µg/µL protamine or 1:12 (v/v) with 0.15 µg/µL RNase 7, to facilitate visualization of the degradation products, for various times points at 37°C. Bacteria were separated from peptide cleavage products by centrifugation, and supernatants were combined with 2× Tricine sample buffer (Beyotime, China) or 5× SDS-PAGE protein loading buffer (Yeasen biotechnology, China), then boiled and frozen at -20°C. Peptide cleavage products were heated at 96°C for 10 min and resolved by 16.5% Tris-Tricine SDS-PAGE (Beyotime, China) or 13% SDS-PAGE. After fixation for 30 min in 5% glutaraldehyde and then washing for 30 min with deionized water, the peptides were stained for 1 h with coomassie blue G-250.

### Fluorescence resonance energy transfer (FRET) activity assay

The synthetic FRET substrate containing ortho-aminobenzoic acid (Abz) as the fluorescent and group 2, 4 nitrophenyl (Dnp) as the quencher and a dibasic motif (RK) in its center (2Abz-SLGRKIQI-K(Dnp)-NH2) was purchased from GL Biochem Ltd. (China) (http://www.glschina.com/en/profile.htm). To perform the assay, bacteria were grown in N-minimal medium to mid-exponential phase and normalized to an OD_600_ nm of 0.6-0.8. Bacterial cells were pelleted by centrifugation, resuspended in PBS (pH 7.4) and normalized to a bacterial density of 3× 10^8^ CFU/mL. Bacteria (∼2.25× 10^7^ CFU in 75 µL) were mixed in a 96-well plate with 75 µL of the FRET substrate (final concentration 40 µM). The fluorescence emission was monitored over 360 min at 25°C with an excitation wavelength at 325 nm and emission wavelength at 430 nm by using a BioTek Synergy 2 plate reader. Initial background measurements were subtracted from the final reaction sample values. Kinetic parameters (*K*_m_, *k*_cat_ and *k*_cat_/*K*_m_) were determined by measuring OmpT activity at 0-240 μM substrate and subsequent fitting of the resulting Michaelis-Menten equation.

### Colorimetric assay

Samples of membrane fractions were diluted in buffer A to appropriate concentrations prior to OmpT activity measurements. OmpT activity was measured in a coupled spectrophotometric assay using the chromogenic substrate IAA-Arg-Arg-pNA purchased from GL Biochem Ltd. (China). Assay conditions in 200 µL reaction system were 100 µg total outer membrane, 0.5 mM IAA-Arg-Arg-pNA, 1 mM Tween 20, 20 mM Mes (pH 7.0) and 0.5 U·mL^-1^ aminopeptidase M. OmpT specific cleavage between the two arginines results in the liberation of Arg-pNA. This compound is then cleaved by aminopeptidase M (Sigma, USA) (being present in excess such that its action is not rate-limiting) resulting in the release of pNA, which is detected spectrophotometrically at 405 nm over 12 h at 37°C by using a BioTek Synergy 2 plate reader. Initial background measurements were subtracted from the final reaction sample values. The above two experimental data were drawn into kinetic curve, and used GraphPard Prim 7 software for differential analysis.

### Inhibition of proteolytic activity

Two serine protease inhibitors (leupeptin [Beyotime, China] and aprotinin [Sigma, USA]) were chose to determine the properties of Asp^267^ (aspartic acid, D) and Ser^276^ (serine, S) of cOmpT by FRET activity assay. Briefly, bacteria were grown in N-minimal medium to mid-exponential phase and normalized to an OD_600_ nm of 0.6-0.8. Bacterial cells were pelleted by centrifugation, resuspended in PBS (pH 7.4) and normalized to a bacterial density of 3× 10^8^ CFU/mL. Assay conditions in 175 µL reaction system in a 96-well plate were bacteria (∼2.25× 10^7^ CFU), 40 μM FRET substrate, 800 μM leupeptin or 400 μM aprotinin. Fluorescence (an excitation of 325 nm and an emission of 430 nm) was monitored over 360 min at 25°C by using a BioTek Synergy 2 plate reader. Initial background measurements were subtracted from the final reaction sample values. The data were drawn into kinetic curve, and used GraphPard Prim 7 software for differential analysis.

## Data availability

The data supporting the findings of the study are available in the article, available upon request from the corresponding author or available from the protein structure data bank PDB (https://www1.rcsb.org/, identifiers 7XW0).

**Expanded View** for this article is available online

## Acknowledgements

This study was supported by grants from the National Key R&D Program of China (2021YFD1800404, 2017YFD0500203-2, 2017YFD0500705), the National Natural Science Foundation of China (31672553, 31972711), the National Program for High Technology Research and Development in China (2003 AA 222141), the Special Fund for Agroscientific Research in the Public Interest (201303044) to S.G., and the Project Funded by the Priority Academic Program Development of Jiangsu Higher Education Institutions (PAPD).

## Author contributions

Gao S., Liu J.H., Wang W.W., and Liu X.F. designed research; Gao Q.Q., Ran T.T. and Huan C.C. provided materials; Liu J.H., Jiang L.Y., Ran T.T. and Wang H. performed research; Liu J.H., Gao S., Wang W.W., Jiang L.Y., and Ran T.T. analyzed data; Liu J.H., Jiang L.Y. and Ran T.T. wrote the paper.

## Conflict of interest

The authors declare that they have no conflict of interest.

## References

Adams PD, Afonine PV, Bunkoczi G, Chen VB, Davis IW, Echols N, Headd JJ, Hung LW, Kapral GJ, Grosse-Kunstleve RW et al (2010) PHENIX: a comprehensive Python-based system for macromolecular structure solution. Acta Crystallogr D Biol Crystallogr 66: 213–221

Brannon JR, Burk DL, Leclerc JM, Thomassin JL, Portt A, Berghuis AM, Gruenheid S, Le Moual H (2015) Inhibition of outer membrane proteases of the omptin family by aprotinin. Infect Immun 83: 2300–2311

Brannon JR, Thomassin JL, Desloges I, Gruenheid S, Le Moual H (2013) Role of uropathogenic Escherichia coli OmpT in the resistance against human cathelicidin LL-37. FEMS Microbiol Lett 345: 64–71

Brannon JR, Thomassin JL, Gruenheid S, Le Moual H (2015) Antimicrobial peptide conformation as a structural determinant of omptin protease specificity. J Bacteriol 197: 3583–3591.

Brogden KA (2005) Antimicrobial peptides: pore formers or metabolic inhibitors in bacteria? Nat Rev Microbiol 3: 238–250

Caulfield AJ, Walker ME, Gielda LM, Lathem WW (2014) The Pla protease of Yersinia pestis degrades fas ligand to manipulate host cell death and inflammation. Cell Host Microbe 15: 424–434

Chen J (2016) Construction of compT, pompT knockout mutants of in avian pathogenic E.coli E058 and evaluation their pathogenicity. Yangzhou university

Dekker N, Cox RC, Kramer RA, Egmond MR (2001) Substrate specificity of the integral membrane protease OmpT determined by spatially addressed peptide libraries. Biochemistry 40: 1694–1701.

Desloges I, Taylor JA, Leclerc JM, Brannon JR, Portt A, Spencer JD, Dewar K, Marczynski GT, Manges A, Gruenheid S et al (2019) Identification and characterization of OmpT-like proteases in uropathogenic Escherichia coli clinical isolates. Microbiologyopen 8: e915

Egile C, D’Hauteville H, Parsot C, Sansonetti PJ (2010) SopA, the outer membrane protease responsible for polar localization of IcsA in Shigella flexneri. Mol Microbiol 23: 1063–1073

Emsley P, Cowtan K (2004) Coot: model-building tools for molecular graphics. Acta Crystallogr D 60: 2126–2132

Evans P (2006) Scaling and assessment of data quality. Acta Crystallogr D 62: 72–82

Franco AA, Kothary MH, Gopinath G, Jarvis KG, Grim CJ, Hu L, Datta AR, McCardell BA, Tall BD (2011) Cpa, the outer membrane protease of Cronobacter sakazakii, activates plasminogen and mediates resistance to serum bactericidal activity. Infect Immun 79: 1578–1587

Gallo RL, Hooper LV (2012) Epithelial antimicrobial defence of the skin and intestine. Nat Rev Immunol 12: 503–516

Gao S, Liu XF, Zhang RK (1996) An improved method for rapid isolation of the outer menbrane proteins from Escherichia coli isolates of chicken origin. Microbiology china, 23: 122–124

Gao S, Zhu XB, Cui HP (1999) The isolation and identification of pathogenic Escherichia coli isolates of chicken origin from some regions in China. Acta Veterinaria et Zootechnica Sinica, 30: 164–171.

Gardell SJ, Craik CS, Hilvert D, Urdea MS, Rutter WJ (1985) Site-directed mutagenesis shows that tyrosine 248 of carboxypeptidase A does not play a crucial role in catalysis. Nature 317: 551–555

Gill RT, Delisa MP, Shiloach M, Holoman TR, Bentley WE (2000) OmpT expression and activity increase in response to recombinant chloramphenicol acetyltransferase overexpression and heat shock in E. coli. J Mol Microb Biotech 2: 283

Grodberg J, Dunn JJ (1988) ompT encodes the Escherichia coli outer membrane protease that cleaves T7 RNA polymerase during purification. J Bacteriol 170: 1245–1253

Grodberg J, Dunn JJ (1989) Comparison of Escherichia coli K-12 outer membrane protease OmpT and Salmonella typhimurium E protein. J Bacteriol 171: 2903–2905

Haiko JM, Suomalainen, Ojala T, Lahteenmaki K, Korhonen TK (2009) Invited review: breaking barriers--attack on innate immune defences by omptin surface proteases of enterobacterial pathogens. Innate Immun 15: 67–80

Hancock R, Sahl HG (2006) Antimicrobial and host-defense peptides as new anti-infective therapeutic strategies. Nat Biotechnol 24: 1551–1557

Hilchie AL, Wuerth K, Hancock RE (2013) Immune modulation by multifaceted cationic host defense (antimicrobial) peptides. Nat Chem Biol 9: 761–768

Hritonenko V, Stathopoulos C (2007) Omptin proteins: an expanding family of outer membrane proteases in Gram-negative Enterobacteriaceae. Mol Membr Biol 24: 395–406

Hui CY, Guo Y, He QS, Liang P, Wu SC, Hong C, Huang SH (2010) Escherichia coli outer membrane protease OmpT confers resistance to urinary cationic peptides. Microbiol Immunol 54: 452–459

Hwang BY, Varadarajan N, Li H, Rodriguez S, Iverson BL, Georgiou G (2007) Substrate specificity of the Escherichia coli outer membrane protease OmpP. J Bacteriol 189: 522–530

Jiang Y, Chen B, Duan CL, Sun BB, Yang JJ, Yang S (2015) Multigene editing in the Escherichia coli genome via the CRISPR-Cas9 system. Appl Environ Microbiol 81: 2506–2514

Johnson TJ, Wannemuehler Y, Johnson SJ, Stell AL, Doetkott C, Johnson JR, Kim KS, Spanjaard L, Nolan LK (2008) Comparison of extraintestinal pathogenic Escherichia coli strains from human and avian sources reveals a mixed subset representing potential zoonotic pathogens. Appl Environ Microbiol 74: 7043–7050

Kabsch W (2010) Integration, scaling, space-group assignment and post-refinement. Acta Crystallogr D 66: 133–144

Korhonen TK, Haiko J, Laakkonen L, Jarvinen HM, Westerlund-Wikstrom B (2013) Fibrinolytic and coagulative activities of Yersinia pestis. Front Cell Infect Microbiol 3: 35

Koten B, Simanski M, Glaser R, Podschun R, Schroder JM, Harder J (2009) RNase 7 contributes to the cutaneous defense against Enterococcus faecium. PLoS One 4: e6424

Kramer RA, Dekker N, Egmond MR (2000) Identification of active site serine and histidine residues in Escherichia coli outer membrane protease OmpT. FEBS Lett 468: 220–224

Kramer RA, Vandeputte-Rutten L, Roon G, Gros P, Dekker N, Egmond MR (2001) Identification of essential acidic residues of outer membrane protease OmpT supports a novel active site. FEBS Lett, 505: 426–430

Kukkonen M, Korhonen TK (2004) The omptin family of enterobacterial surface proteases/adhesins: from housekeeping in Escherichia coli to systemic spread of Yersinia pestis. Int J Med Microbiol 294: 7–14

Kukkonen M, Lahteenmaki K, Suomalainen M, Kalkkinen N, Emody L, Lang H, Korhonen TK (2001) Protein regions important for plasminogen activation and inactivation of alpha2-antiplasmin in the surface protease Pla of Yersinia pestis. Mol Microbiol 40: 1097–1111

Lathem WW, Price PA, Miller VL, Goldman WE (2007) A plasminogen-activating protease specifically controls the development of primary pneumonic plague. Science 315: 509–513

Le Sage V, Zhu L, Lepage C, Portt A, Viau C, Daigle F, Gruenheid S, Le Moual H (2009) An outer membrane protease of the omptin family prevents activation of the Citrobacter rodentium PhoPQ two-component system by antimicrobial peptides. Mol Microbiol 74: 98–111

Liu J, Mu X, Wang X, Huan H, Gao Q, Chen J, Qiao P, Jiang L, Gao S, Liu X (2019) Unexpected transcriptome pompT’ contributes to the increased pathogenicity of a pompT mutant of avian pathogenic Escherichia coli. Vet Microbiol 228: 61–68

McCarter JD, Stephens D, Shoemaker K, Rosenberg S, Kirsch JF, Georgiou G (2004) Substrate specificity of the Escherichia coli outer membrane protease OmpT. J Bacteriol 186: 5919–5925

Mccoy AJ, Grosse-Kunstleve RW, Adams PD, Winn MD, Storoni LC, Read RJ (2007) Phaser crystallographic software. J Appl Crystallogr 40: 658–674

Mcphee JB, Small CL, Reid-Yu SA, Brannon JR, Moual HL, Coombes BK (2014) Host defense peptide resistance contributes to colonization and maximal intestinal pathology by Crohn’s disease-associated adherent-invasive Escherichia coli. Infect Immun 82: 3383–3393

Murshudov GN, Vagin AA, Dodson EJ (1997) Refinement of macromolecular structures by the maximum-likelihood method. Acta Crystallogr D Biol Crystallogr 53: 240–255

Nelson DL, Kennedy EP (1971) Magnesium transport in Escherichia coli: inhibition by cobaltous ion. J Biol Chem 246: 3042–3049

Piers KL, Hancock RE (1994) The interaction of a recombinant cecropin/melittin hybrid peptide with the outer membrane of Pseudomonas aeruginosa. Mol Microbiol 12: 951–958

Samantha G, Le MH (2012) Resistance to antimicrobial peptides in Gram-negative bacteria. FEMS Microbiol Lett 2: 81–89

Sodeinde OA, Goguen JD (1989) Nucleotide sequence of the plasminogen activator gene of Yersinia pestis: relationship to ompT of Escherichia coli and gene E of Salmonella typhimurium. Infect Immun 57: 1517–1523

Sodeinde OA, Subrahmanyam YV, Stark K, Quan T, Bao Y, Goguen JD (1992) A surface protease and the invasive character of plague. Science 258: 1004–1007

Stathopoulos C (1998) Structural features, physiological roles, and biotechnological applications of the membrane proteases of the OmpT bacterial endopeptidase family: A micro-review. Membr cell biol 12: 1–8

Stumpe S, Schmid R, Stephens DL, Georgiou G, Bakker EP (1998) Identification of OmpT as the protease that hydrolyzes the antimicrobial peptide protamine before it enters growing cells of Escherichia coli. J Bacteriol 180: 4002–4006

Sugimura K, Higashi N (1988) A novel outer-membrane-associated protease in Escherichia coli. J Bacteriol 170: 3650–3654

Sugimura K, Nishihara T (1988) Purification, characterization, and primary structure of Escherichia coli protease VII with specificity for paired basic residues: identity of protease VII and OmpT. J Bacteriol 170: 5625

Thomassin JL, Brannon JR, Gibbs BF, Gruenheid S, Le Moual H (2012) OmpT outer membrane proteases of enterohemorrhagic and enteropathogenic Escherichia coli contribute differently to the degradation of human LL-37. Infect Immun 80: 483–492

Thomassin JL, Brannon JR, Kaiser J, Gruenheid S, Le Moual H (2012) Enterohemorrhagic and enteropathogenic Escherichia coli evolved different strategies to resist antimicrobial peptides. Gut Microbes 3: 556–561

Vandeputte-Rutten L, Kramer RA, Kroon J, Dekker N, Egmond MR, Gros P (2001) Crystal structure of the outer membrane protease OmpT from Escherichia coli suggests a novel catalytic site. EMBO J 20: 5033–5039

Varadarajan N, Gam J, Olsen MJ, Georgiou G, Iverson BL (2005) Engineering of protease variants exhibiting high catalytic activity and exquisite substrate selectivity. P Natl Acad Sci USA 102: p.6855-6860

Wang H, Schwaderer AL, Kline J, Spencer JD, Kline D, Hains DS (2013) Contribution of structural domains to the activity of ribonuclease 7 against uropathogenic bacteria. Antimicrob Agents Chemother 57: 766–774

He XL, Wang Q, Peng L, Qu YR, Santhosh (2015) Role of uropathogenic Escherichia coli outer membrane protein T in pathogenesis of urinary tract infection. Pathog Dis, 73: ftv006

Yam CH, Siu WY, Kaganovich D (2001) Cleavage of cyclin A at R70/R71 by the bacterial protease OmpT. P Natl Acad Sci USA 98: 497–501

Zasloff, Michael (2002) Antimicrobial peptides of multicellular organisms. Nature 415: 389–395

Zhang L, Rozek A, Hancock RE (2001) Interaction of cationic antimicrobial peptides with model membranes. J Biol Chem 276: 35714–35722

